# Loss of PRMT5 promotes PDGFRα degradation during oligodendrocyte differentiation and myelination

**DOI:** 10.1101/252056

**Authors:** Sara Calabretta, Gillian Vogel, Zhenbao Yu, Karine Choquet, Lama Darbelli, Thomas B. Nicholson, Claudia L. Kleinman, Stéphane Richard

**Affiliations:** Segal Cancer Center, Bloomfield Center for Research on Aging, Lady Davis Institute for Medical Research and Departments of Oncology and Medicine, McGill University, Montréal, Québec, Canada H3T 1E2; Segal Cancer Centre, Lady Davis Institute for Medical Research, Sir Mortimer B. Davis Jewish General Hospital, and Department of Human Genetics, Faculty of Medicine, McGill University, Montréal, Québec, Canada H3T 1E2; Developmental and Molecular Pathways to Chemical Biology and Therapeutics, Novartis Institute for Biomedical Research, 250 Massachusetts Avenue, Cambridge, Massachusetts 02139, USA

**Keywords:** Arginine methylation, PRMT5, PDGFRα, Cbl, myelination, cellular differentiation

## Abstract

Platelet derived growth factor receptor α (PDGFRα) signaling is required for proliferation, commitment and maintenance of oligodendrocyte (OL) precursor cells (OPCs). PDGFRα signaling promotes OPC homeostasis and its attenuation signals OPC differentiation and maturation triggering the onset of myelination of the central nervous system (CNS). The initial steps of how PDGFRα signaling is attenuated are still poorly understood. Herein we show that decreased Protein Arginine MethylTransferase5 (PRMT5) expression, as occurs during OPC differentiation, is involved in the down-regulation of PDGFRα by modulating its cell surface bioavailability leading to its degradation in a Cbl-dependent manner. Mechanistically, loss of arginine methylation at R554 of the PDGFRα intracellular domain reveals a masked Cbl binding site at Y555. Physiologically, depletion of PRMT5 in OPCs results in severe CNS myelination defects. We propose that decreased PRMT5 activity initiates PDGFRα degradation to promote OL differentiation. More broadly, inhibition of PRMT5 may be used therapeutically to manipulate PDGFRα bioavailability.

## Introduction

Arginine methylation is a common post-translational modification that impacts multiple biological processes from gene expression, mRNA splicing, DNA damage and signal transduction (Blanc and Richard, 2017). Three types of methylated arginine residues are observed: ω–*N^G^* –monomethylarginine (MMA), *ω-N^G^, N^G^* –asymmetric dimethylarginine (aDMA), and *ω–N^G^, N’^G^* –symmetric dimethylarginine (sDMA) (Bedford and Clarke, 2009). In mammalian cells, the family of enzymes responsible for the transferring of methyl groups from S-adenosyl-methionine to the nitrogen atoms of arginine are called Protein Arginine Methyltransferases (PRMTs). The major type II enzyme catalysing sDMA is PRMT5 and it exists as an octameric complex with its cofactor, methylosome protein 50 (MEP50/WDR77) (Antonysamy et al., 2012; Friesen et al., 2001).

PRMT5 is a well-known transcriptional co-repressor that methylates histone H3R8 and H4R3 to repress gene expression (Pal et al., 2004; Zhao et al., 2009). PRMT5 also has numerous non-histone related functions: PRMT5 directly methylates transcription factors E2F1 and p53 to regulate their function (Cho et al., 2012; Jansson et al., 2008). Moreover, PRMT5 regulates small nuclear ribonucleoprotein complex biogenesis by methylating Sm proteins (Friesen et al., 2001) and as such defects in pre-mRNA splicing containing weak 5’ donor sites are observed in PRMT5-deficient neural progenitor cells (Bezzi et al., 2013). PRMT5-mediated maintenance of splicing fidelity was also shown in cancer, particularly, in lymphoma where it was identified as a component of Myc-mediated tumorigenesis (Koh et al., 2015). PRMT5 and sDMA have been implicated in tumorigenesis. PRMT5 cooperates with the oncogenic mutant of CyclinD1 (D1T286A) in dampening p53 activity in response to DNA damage and promotes the activity of c-MYC, NOTCH1, and MLL-AF9 oncogenes (Kaushik et al., 2017; Li et al., 2015). Although the role of PRMT5 in regulating splicing and transcription is well-defined, little is known of its cytoplasmic role in receptor signaling, however, PRMT5 methylates the epidermal growth factor receptor (EGFR) and RAF proteins to dampen the RAS-ERK pathway (Andreu-Perez et al., 2011; Hsu et al., 2011).

PDGFRα is a receptor tyrosine kinase (RTK) that modulates several aspects of oligodendrocyte progenitor cells (OPCs) biology, such as proliferation, migration and commitment to oligodendrocytes (OLs) (Armstrong et al., 1990; Barres et al., 1992; Calver et al., 1998; Fruttiger et al., 1999). PDGFRα signaling is activated by the binding of its ligand PDGF-AA secreted in the central nervous system (CNS) by astrocytes and neurons (Raff et al., 1988; Richardson et al., 1988; Yeh et al., 1991). PDGFRα haploinsufficiency causes acceleration of OPC maturation with altered OL differentiation timing and migration (McKinnon et al., 2005). PDGF-A knockout mice that survive up to the third post-natal week display a shaking phenotype, accompanied by reduced numbers of OPCs in the CNS and hypomyelination (Fruttiger et al., 1999). PDGFRα engages several signaling pathways such as the RAS-ERK pathway, which is a positive stimulus for OPC differentiation and CNS myelination (Fyffe-Maricich et al., 2011; Ishii et al., 2012; Xiao et al., 2012). PDGFRα activity is fine-tuned through several mechanisms, including protein degradation by the Cbl E3 ligase that triggers ubiquitination promoting degradation (Miyake et al., 1998).

We have previously shown that PRMT5-deficiency impairs OL differentiation *in vitro* (Huang et al., 2011). It is known that PDGFRα expression decreases during differentiation to allow OL maturation (Hart et al., 1989), however, the early steps of this process are not entirely understood. Herein we show that PRMT5 deficiency regulates early steps of OL differentiation *in vivo*. We also show that PRMT5 regulates PDGFRα during OL differentiation by orchestrating its degradation in a Cbl-dependent manner, masking a novel binding site through arginine methylation. Our findings suggest that the manipulation of PRMT5 levels and activity may be used in a broader sense to dampen PDGFRα signaling.

## Results

### PRMT5 expression decreases during oligodendrocyte differentiation

PRMT5 expression is required for proper brain development, as a conditional knockout mouse model for CNS depletion of PRMT5 (Nestin-CRE) caused early post-natal mortality and defects in brain development (Bezzi et al., 2013). PRMT5 immunoblots of total mouse brain lysates give rise to two distinct bands (Huang et al., 2011) and both disappear when preabsorbed with the immunogenic peptide (Figure 1A). Although the identity and the cell origin of the upper band is unknown, the lower band decreases during development correlating with decreased PRMT5 and WDR77 mRNA levels (Figure 1B) and overall decrease in proteins harboring sDMA, as assessed by anti-sDMA antibodies (Figure 1A). The lower band corresponds to PRMT5 and is observed in E14.5 embryos and OPCs (Figure 1C). Furthermore, the lower PRMT5 band decreases during OL differentiation, as does the mRNAs for PRMT5 and WDR77 (Figure 1D, 1E) and is the band that decreases in the *PRMT5^FL/FL;Olig2Cre^* mice (see below; Figure S1C). Taken together these findings suggest that during OL differentiation, PRMT5 expression and activity decreases.

**Figure 1.**
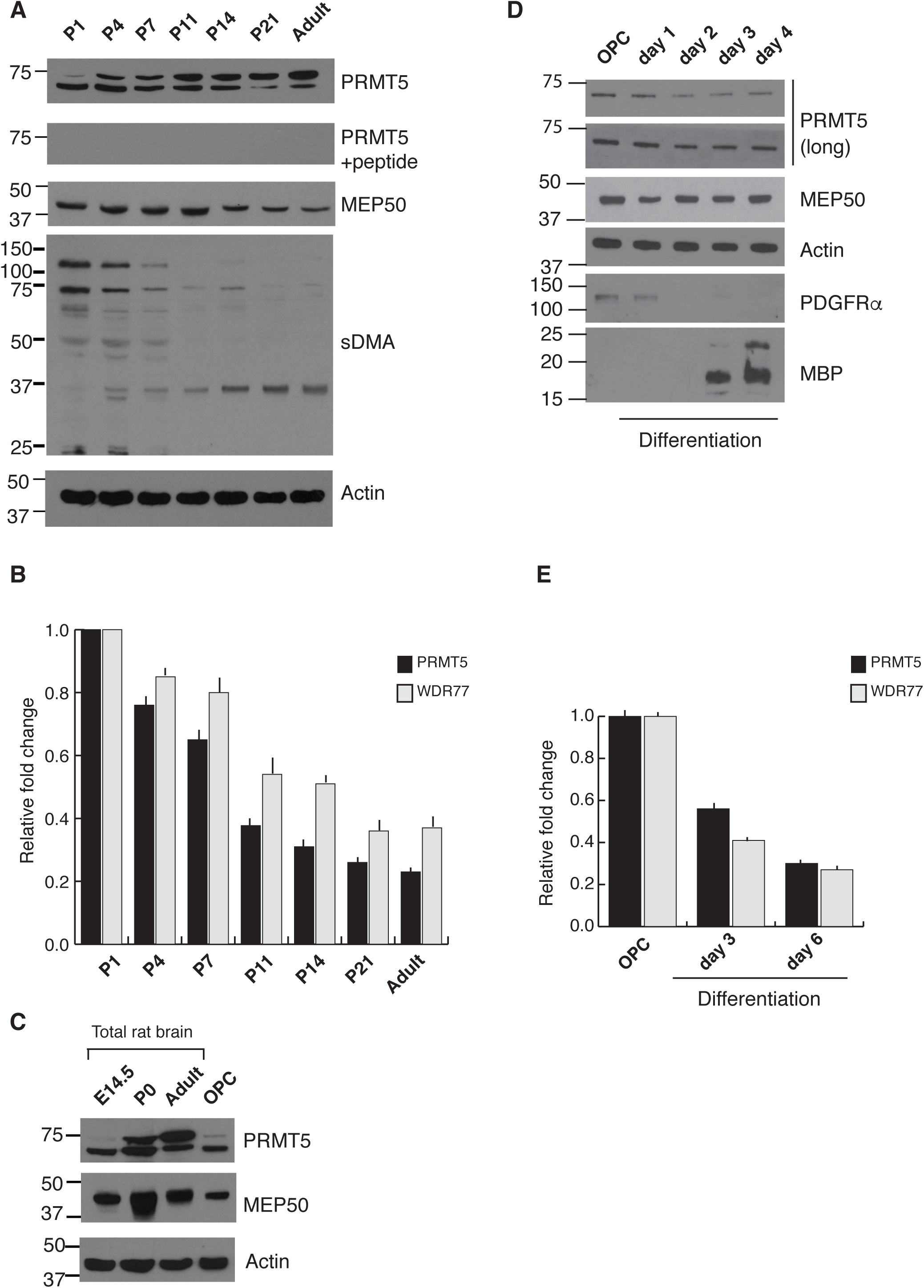
PRMT5 expression decreases during OL differentiation and during brain development. **A.** Total brain lysates from post-natal day 1 (P1), P4, P7, P11, P14, P21 and from adult mice were prepared and separated by SDS-PAGE. Immunoblotting analysis of PRMT5 with anti-PRMT5 antibody in the presence or absence of the immunogenic peptide, MEP50 and symmetrical methylation (sDMA) was performed. β-actin was used as loading control. Molecular mass markers are on the left in kDa. **B.** RT-qPCR analysis of PRMT5 and WDR77 mRNAs was performed using total RNA isolated from the indicated mouse brains. Values were normalized to 18S and GADPH mRNAs. Each bar represents the mean fold change ± S.D. **C.** Western blot analysis of PRMT5 and MEP50 in total rat brain of the indicated age and primary rat OPCs. β-actin was used as loading control. Molecular mass markers are on the left in kDa. **D.** Western blot analysis of PRMT5 and MEP50 during OPC differentiation. β-actin was used as loading control. PDGFRα and MBP were used to assess OPCs and mature OLs, respectively. Molecular mass markers are on the left in kDa. **E.** RT-qPCR analysis of PRMT5 and WDR77 mRNA expression in rat OPCs and differentiated OLs. Values were normalized to mRNA levels of HPRT and GAPDH. Bar graphs represent the mean fold change ± S.D.

### OL-specific knockout of PRMT5 results in dysmyelination and early post-natal death

Methylation by PRMT5 is a known regulator of cellular differentiation (Bezzi et al., 2013; Tee et al., 2010), however, whether it regulates CNS myelination remains undefined. To investigate the role of PRMT5 *in vivo*, we generated PRMT5-deficient OLs using the Olig2-Cre driver (Figure S1A) (Schuller et al., 2008). Recombination at the *Prmt5* locus was confirmed by genomic PCR, as well as reduced PRMT5 mRNA and protein levels (Figure S1B-E). *PRMT5^FL/FL;Olig2Cre^* mice were born at the expected Mendelian ratio and began displaying rapid tremors by post-natal day 10 followed by death prior to the third post-natal week (Figure 2A), resembling a dysmyelination mouse model like the *qkI* mice (Darbelli et al., 2016). We next analysed the expression of the myelin basic proteins (MBP) by immunohistochemical analysis of post-natal day 14 (P14) coronal brain sections. We observed very little anti-MBP staining within the *corpus callosum* of the *PRMT5^FL/FL;Olig2Cre^* mice, suggesting a lack of mature OLs and myelination defects (Figure 2B). The absence of mature compacted myelin was further confirmed by the diminished fluoromyelin staining in the *PRMT5^FL/FL;Olig2Cre^* mice (Figure 2C). Moreover, *PRMT5^FL/FL;Olig2Cre^* mice showed decreased levels of several markers of mature OLs including MBP, myelin oligodendrocyte glycoprotein (MOG), myelin-associated glycoprotein (MAG), 2’,3’-Cyclic-nucleotide 3’-phosphodiesterase (CNP) and proteolipid protein (PLP) at both the mRNA and protein levels in whole brains (Figure 2D, 2E). However, no profound changes were observed in the expression of glial fibrillary acidic protein (GFAP), a marker of astrocytes, nor in the expression of neurofilament light polypeptide (Nefl), a marker of neurons (Figure 2D, 2E). These data demonstrate that conditional loss of PRMT5 in the OL lineage leads to hypomyelination within the CNS resulting in early post-natal death.

**Figure 2.**
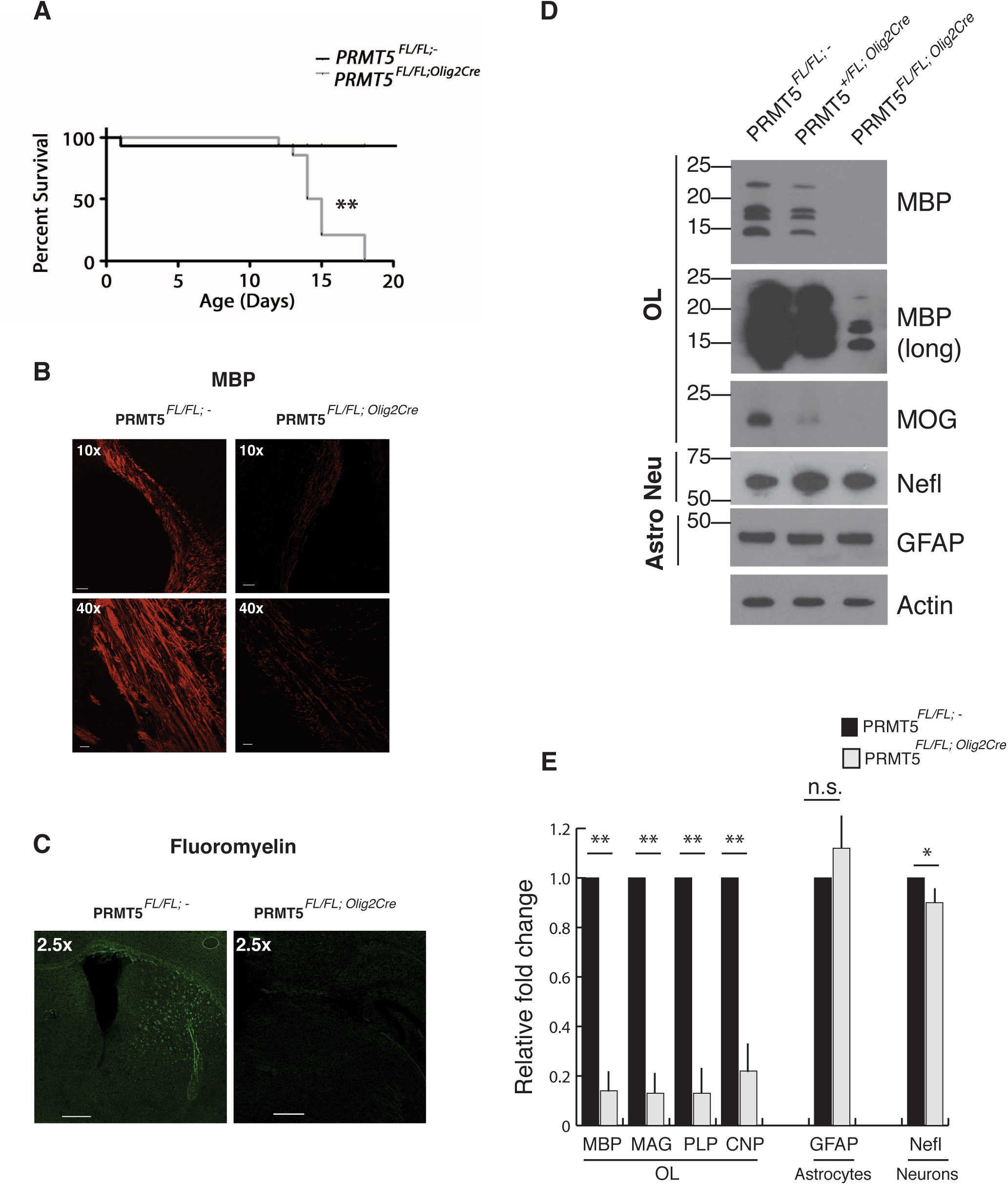
PRMT5 conditional depletion causes hypomyelination and death by the third post-natal week. **A.** Kaplan-Meier survival curve of *PRMT5^FL/FL:-^* (n=15, black) and *PRMT5^FL/FL;Olig2Cre^* (n=15, gray) mice. *p* value was determined by Log-rank test (** *p* ≤ 0.01). **B.** Representative images of coronal sections of P14 brains of *PRMT5^FL/FL:-^* and *PRMT5^FL/FL;Olig2Cre^* mice stained with anti-MBP antibody at the indicated magnification (n=3 mice per genotype). The area shown is the corpus callosum. Scale bars: 50 μm for 10x and 10 μm for 40x. **C.** Representative images of coronal sections from P14 brains of *PRMT5^FL/FL:-^* and *PRMT5^FL/FL;Olig2Cre^* mice stained with fluoromyelin at 2.5x magnification (n=2 mice per genotype). Scale bar: 400 μm. **D. E.** Western blot and RT-qPCR analysis of brain lineage markers in *PRMT5^FL/FL:-^*, *PRMT5^FL/+:Olig2-Cre^* and *PRMT5^FL/FL:Olig2-Cre^* P14 mice. MBP, MOG, MAG, PLP, and CNP protein and mRNA expression was analysed as markers of mature OLs. Nefl protein and mRNA expression was analysed as a neuronal marker (Neu) and GFAP protein and mRNA expression as a marker of astrocytes (Astro). β-actin was used as an immunoblotting loading control. RT-qPCR values were normalized to GAPDH mRNA. Bar graphs represent the mean fold change ± S.D. (n=3). *p* value was determined by two-tailed t-test (\**p* ≤ 0.05; \*\**p* ≤ 0.01; n.s. = not significant).

### PRMT5 deficiency blocks oligodendrocytes in the precursor stage

To investigate whether the lack of myelin of *PRMT5^FL/FL;Olig2Cre^* mice was a result of a complete loss of the OL lineage or due to a block at a certain stage of OL differentiation, we analysed the expression of OL markers by immunofluorescence in the *corpus callosum* of P14 coronal sections. We analysed Olig2 expression as a pan-OL marker and we observed a significant decrease in the number of Olig2 positive cells in *PRMT5^FL/FL;Olig2Cre^* mice, suggesting an impairment of a specific stage of OL maturation (Figure 3A). We analysed PDGFRα levels to identify cells of the OPC stage and CC1^+^ as a marker of fully mature OLs. A slight but not significant, decrease of PDGFRα positive cells was observed between *PRMT5^FL/FL:-^* and *PRMT5^FL/FL;Olig2Cre^* mice (Figure 3B), while a highly significant decrease was observed for the CC1^+^ positive cells (Figure 3C). These findings suggest that the hypomyelination observed in the *PRMT5^FL/FL;Olig2Cre^* mice is a consequence of the lack of mature and functional OLs within the brain.

**Figure 3.**
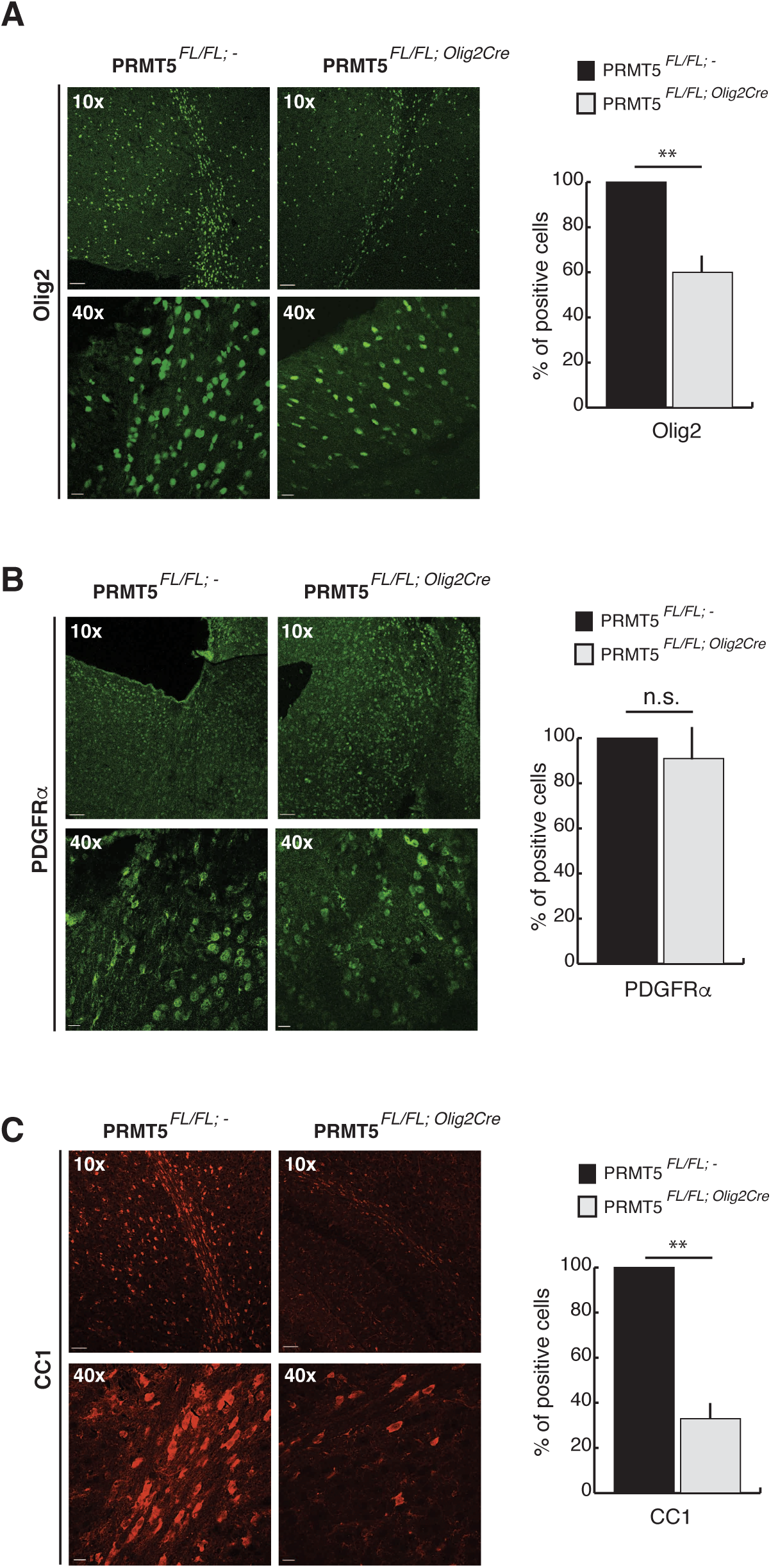
*PRMT5^FL/FL:Olig2-Cre^* hypomyelination is due to loss of mature OLs. **A.B.C.** Representative images of coronal sections of P14 brains of *PRMT5^FL/FL:-^* and *PRMT5^FL/FL;Olig2Cre^* mice at the indicated magnification. The area shown is the corpus callosum. Olig2 staining was analysed as marker of OL lineage, PDGFRα, and CC1 stainings were analysed as marker of precursor and mature OLs, respectively. Bar graphs represent the mean percentage ± S.D. of the indicated positive cells in *PRMT5^FL/FL:-^* (black, set to 100) and *PRMT5^FL/FL;Olig2Cre^* (gray) brain sections (n=3 mice per genotype). *p* value was determined by two-tailed t-test (** *p* ≤ 0.01; n.s. = not significant).

To investigate how PRMT5 depletion in the OL lineage affects the brain transcriptome, we performed RNA sequencing (RNA-seq) analysis to profile mRNA abundance in *PRMT5^FL/FL:-^* and *PRMT5^FL/FL;Olig2Cre^* P14 mouse brains. We performed unsupervised clustering analysis of samples based on expression profiles. With the 1,000 most variant genes, *PRMT5^FL/FL:-^* and *PRMT5^FL/FL;Olig2Cre^* mice formed distinct, robust clusters (Figure S2A-B). Principal Component Analysis (PCA) based on gene expression data show that the first component (PC1) separates the two groups clearly, independently of the number of genes used (Figure S2B). This implies that OL-specific depletion of *PRMT5* causes important changes in gene expression in the brain. Indeed, differential expression analysis showed statistically significant differences in 321 genes, (Figure S2C, Table S1) and many of the top downregulated genes in brains of *PRMT5^FL/FL;Olig2Cre^* mice are involved in the differentiation of OLs and in myelination consistent with myelination defects (i.e. Myrf, Mog, Mag, Mobp, Plp1, Mbp, Cnp; Figure S2C). Gene Ontology (GO) analysis of the most robustly deregulated genes (absolute fold change > 2 and statistically significant) revealed that genes involved in myelination and CNS homeostasis were overrepresented among the downregulated genes, suggesting that these are the principal functions affected in the *PRMT5^FL/FL;Olig2Cre^* mice (Figure S2D). Considering the well-established role of PRMT5 as regulator of splicing fidelity (Bezzi et al., 2013), we analysed the impact of OL-specific PRMT5 depletion on genome-wide exon usage and alternative splicing (AS) methods to infer splicing differences. We observed a total of 265 candidate AS events differentially modulated in the *PRMT5^FL/FL;Olig2Cre^* mice (Figure S2E, Table S1). GO analysis of the alternatively spliced genes revealed that processes involved in CNS homeostasis and myelination were still the principal pathways affected (Figure S2F). Thus, lack of PRMT5 in the OL lineage affects both gene expression and AS of myelin and CNS-specific processes, supporting its well-known role as a gene expression regulator of the OL lineage *in vivo*.

### PRMT5 regulates PDGFRα membrane availability

Morphological analysis of the OPCs was performed in the *corpus callosum* of P14 coronal brain sections of the *PRMT5^FL/FL;Olig2Cre^* mice using an antibody that recognizes an epitope of the extracellular domain of PDGFRα. These experiments revealed that in *PRMT5^FL/FL:-^* mice,

PDGFRα was mainly distributed at the plasma membrane, as expected, while intracellular accumulation and mis-localization of the receptor was mainly observed in the OPCs of *PRMT5^FL/FL;Olig2Cre^* mice (Figure 4A). Accordingly, PDGFRα intensity analysis revealed a decreased perimeter *vs* area ratio in the *PRMT5^FL/FL;Olig2Cre^* OPCs (Figure 4B). Furthermore, the observed altered distribution of PDGFRα was accompanied with decreased PDGFRα protein levels in P14 whole brain extracts of *PRMT5^FL/FL;Olig2Cre^* mice compared to *PRMT5^FL/FL:-^* mice (Figure 4C). Notably, no significant change in PDGFRα mRNA levels was observed in *PRMT5^FL/FL;Olig2Cre^* mice using RNA-seq (Table S1), thus pointing out a role for PRMT5 which goes beyond epigenetic regulation. Collectively, these data suggest a function for PRMT5 in the regulation PDGFRα availability at the plasma membrane and protein stability.

**Figure 4.**
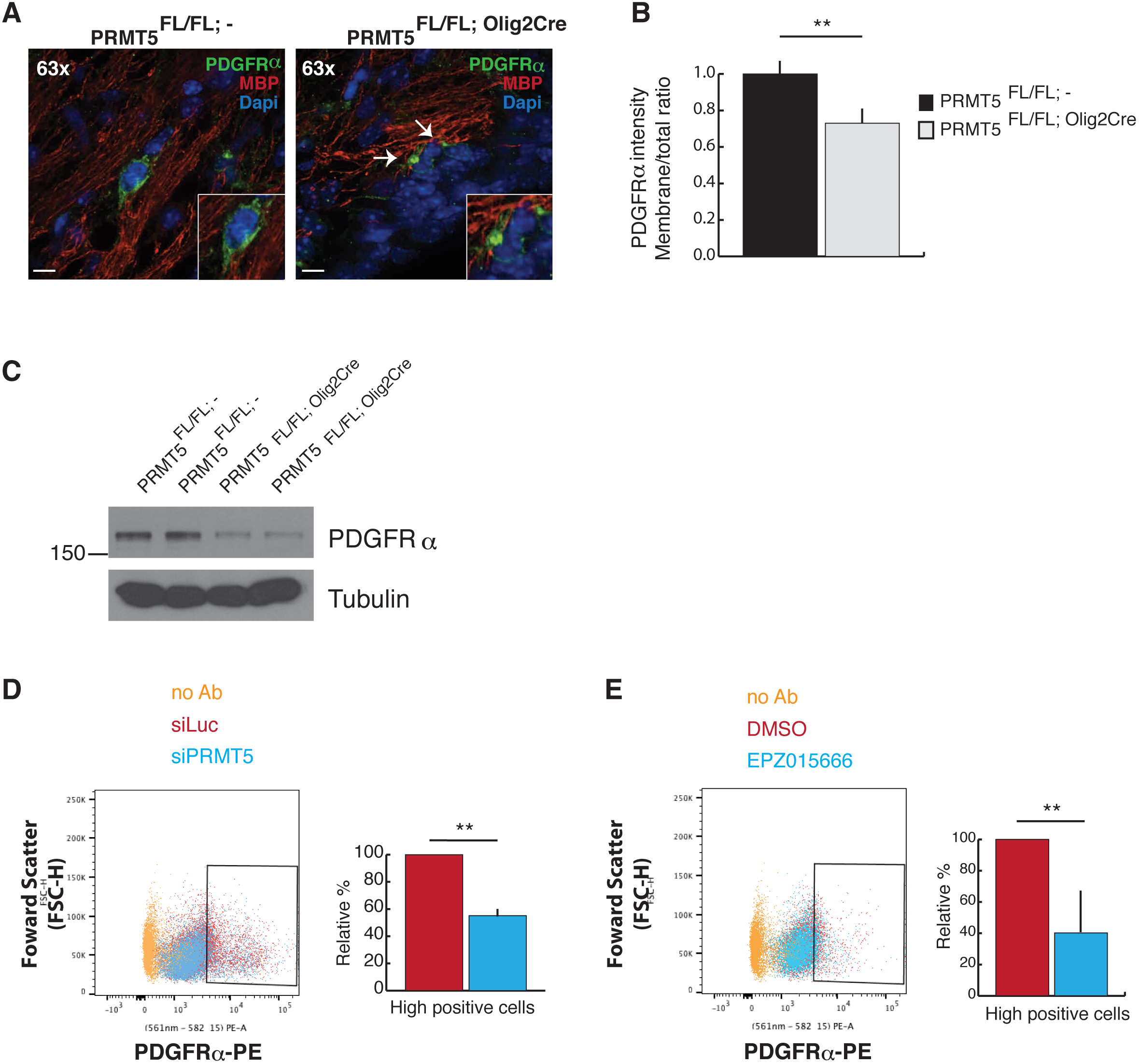
PDGFRα bioavailability requires PRMT5 activity in primary OPCs. **A.** Representative images of coronal sections of P14 brains of *PRMT5^FL/FL:-^* and *PRMT5^FL/FL;Olig2Cre^* mice stained with anti-PDGFRα and anti-MBP antibodies as well as DAPI to visualize nuclei at 63x magnification. The area shown is the corpus callosum and the scale bar represents 4 μm. **B.** Quantification of anti-PDGFRα intensity (perimeter versus area ratio, n=3 mice per genotype, arbitrary units). *PRMT5^FL/FL^*^:-^ ratio set to 1. *p* value was determined by two-tailed ttest (\*\**p* ≤ 0.01). **C.** Western blot analysis of PDGFRα expression in total brain lysates isolated from P14 *PRMT5^FL/FL:-^* and *PRMT5^FL/FL;Olig2Cre^* mice. Tubulin was used as loading control. **D. E.** Representative flow cytometry analysis of rat primary OPC transfected with either siLuciferase or siPRMT5. OPCs were either treated with DMSO or EPZ015666 as indicated. Cells were analysed for the extracellular expression of PDGFRα. PDGFRα high cells were gated as indicated and relative percentage of cells are shown, siLuc and DMSO (red bars) were set as 100%. PRMT5 depletion and inhibition led to decreased cell percentage (blue bars). *p* value was determined by two-tailed t-test (n=3; ** *p* ≤ 0.01).

We next analysed PDGFRα membrane localization by flow cytometry analysis in rat primary OPCs transfected with control siLuc or siPRMT5. We observed a significant decrease of PDGFRα at the plasma membrane in PRMT5-depleted OPCs compared to siLuc cells (Figure 4D). Similar findings were observed, when we impaired PRMT5 activity in OPCs using the PRMT5 inhibitor, EPZ015666 (Figure 4E). These findings indicate that PRMT5 methylation activity is required for the localization of PDGFRα at the cell membrane in OPCs. PDGFRα signaling was next examined in the presence or absence of PRMT5. OPCs treated with DMSO or with EPZ015666 inhibitor were serum-starved for 2 h, then challenged with PDGF-AA at the indicated times. OPCs treated with EPZ015666 showed dampened AKT and ERK activation compared to DMSO control (Figure S3). These findings showed that PDGFRα membrane availability and decreased protein expression resulted in decreased downstream signaling activation. Taken together, these data show that PRMT5 activity is required to ensure PDGFRα membrane bioavailability for downstream signaling.

### PRMT5 regulates PDGFRα protein stability in Cbl-dependent manner

We noted a significant decrease in PDGFRα protein expression in whole brain extracts of *PRMT5^FL/FL;Olig2Cre^* mice (Figure 4C). These findings suggested that PRMT5 mediated accumulation of intracellular PDGFRα led to its degradation. Thus, we investigated the impact of PRMT5 depletion on PDGFRα protein stability. We transfected HEK293T cells with PDGFRα and either siLuc or siPRMT5 and measured protein stability using cycloheximide (CHX) treatment followed by immunoblotting with anti-PDGFRα antibody. PRMT5 depletion decreased PDGFRα protein half-life to ∼ 120 min following CHX treatment while it is >180 min in wild type cells (Figure 5A). This effect was abolished by cotreating cells MG132, but not chloroquine, suggesting the involvement of the proteasome and not the lysosomal compartment (Figure 5A).

**Figure 5.**
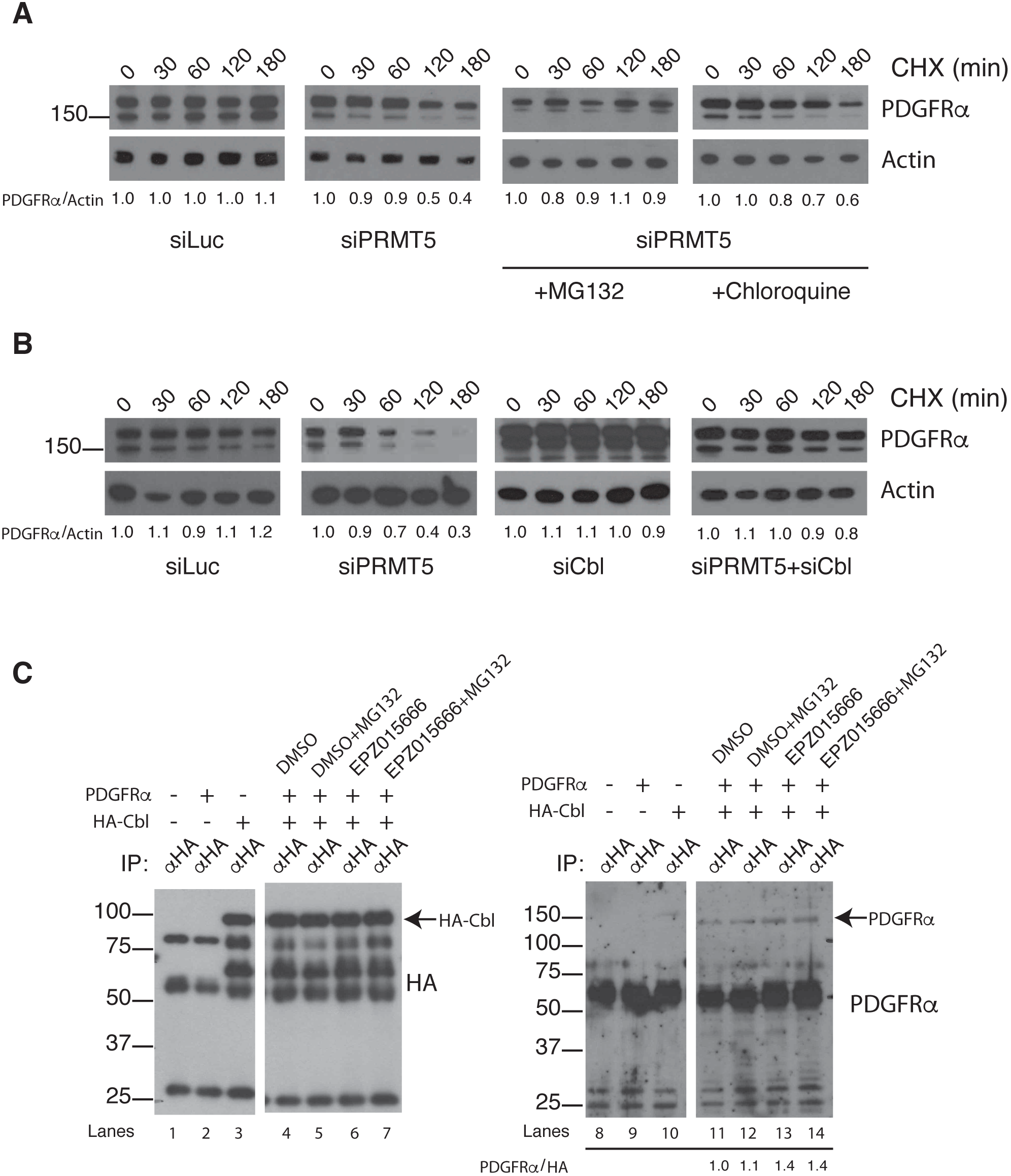
PRMT5 regulate PDGFRα protein stability in a Cbl-dependent manner. **A.** HEK293T cells were transfected an expression vector encoding the PDGFRα along with siLuc (control) or siPRMT5. The PDGFRα/siPRMT5 transfected cells were treated with MG132 or chloroquine. The transfected cells were treated with cycloheximide (CHX) for the indicated time and the cells lysed and immunoblotted with anti-PDGFRα antibodies. β-actin was used as loading control. The relative PDGFRα/Actin ratio was calculated by densitometry and time 0 was set as 1.0. A representative experiment is shown (n=3). **B.** HEK293T cells were transfected an expression vector encoding the PDGFRα along with siLuc, siPRMT5, siCbl, or siPRMT5 + siCbl. The transfected cells were treated with CHX and analysed as in panel A (n=4). **C.** HEK293T transfected with expression vectors encoding HA-Cbl and/or PDGFRα were left untreated or treated with MG132 and/or EPZ015666 as indicated. The cells were lysed and immunoprecipitated with anti-HA antibodies. The bound proteins were either immunoblotted with anti-HA antibodies to detect HA-Cbl (lanes 1-7) or with anti-PDGFRα antibodies (lanes 8-14). The PDGFRα/HA-Cbl ratio is shown, as assessed by densitometric analysis (lanes 8-14). Molecular mass markers are on the left in kDa (n > 5).

A well-established regulator of PDGFRα protein degradation is the Cbl E3 ligase that dampens receptor protein tyrosine kinase (RTK) signaling by promoting their ubiquitination and degradation (Miyake et al., 1998). To investigate the involvement of Cbl in decreased PDGFRα protein stability in the absence of PRMT5, we assessed PDGFRα protein half-life upon CHX treatment in HEK293T transfected with both siPRMT5 and siCbl. Strikingly, the co-depletion of Cbl with PRMT5 stabilized PDGFRα protein levels (Figure 5B), suggesting that PRMT5 depleted cells require Cbl to promote PDGFRα degradation. We next hypothesized that PRMT5 might be involved in the recruitment of Cbl on PDGFRα. We assessed Cbl/PDGFRα interaction by co-immunoprecipitation analysis. HEK293T transfected with HA-Cbl or PDGFRα alone or with both proteins in combination and treated with either DMSO or with EPZ015666, alone or in combination with MG132 (Figure 5C), were lysed and immunoprecipitated using anti-HA antibodies and the bound proteins immunoblotted for PDGFRα. Importantly, EPZ015666 or EPZ015666 + MG132 treatment increased Cbl/PDGFRα interaction with respect to DMSO control (Figure 5C, compare lanes 11 with 13 and 14). Taken together, these data unveil a new role for PRMT5 as a regulator of Cbl binding on PDGFRα.

### PDGFRα is arginine methylated by PRMT5

We investigated whether PDGFRα and PRMT5 interact *in vivo*. We transfected HEK293T cells with expression vectors encoding myc-PRMT5, PDGFRα alone or in combination. Cells were lysed and immunoprecipitated (IP) with anti-Myc antibodies and the bound proteins separated by SDS-PAGE and immunoblotted with anti-PDGFRα antibodies. Indeed, the PDGFRα co-immunoprecipitated with myc-PRMT5 and this was not observed in the control lanes (Figure 6A, upper panel, Figure 6B). We next investigated whether PDGFRα could be a substrate of PRMT5. PDGFRα was expressed with or without myc-PRMT5, and cell lysates were prepared and immunoblotted with anti- PDGFRα antibodies and with anti-sDMA antibodies (Figure 6C). Indeed the PDGFRα was observed to contain sDMA in the presence of PRMT5.

**Figure 6.**
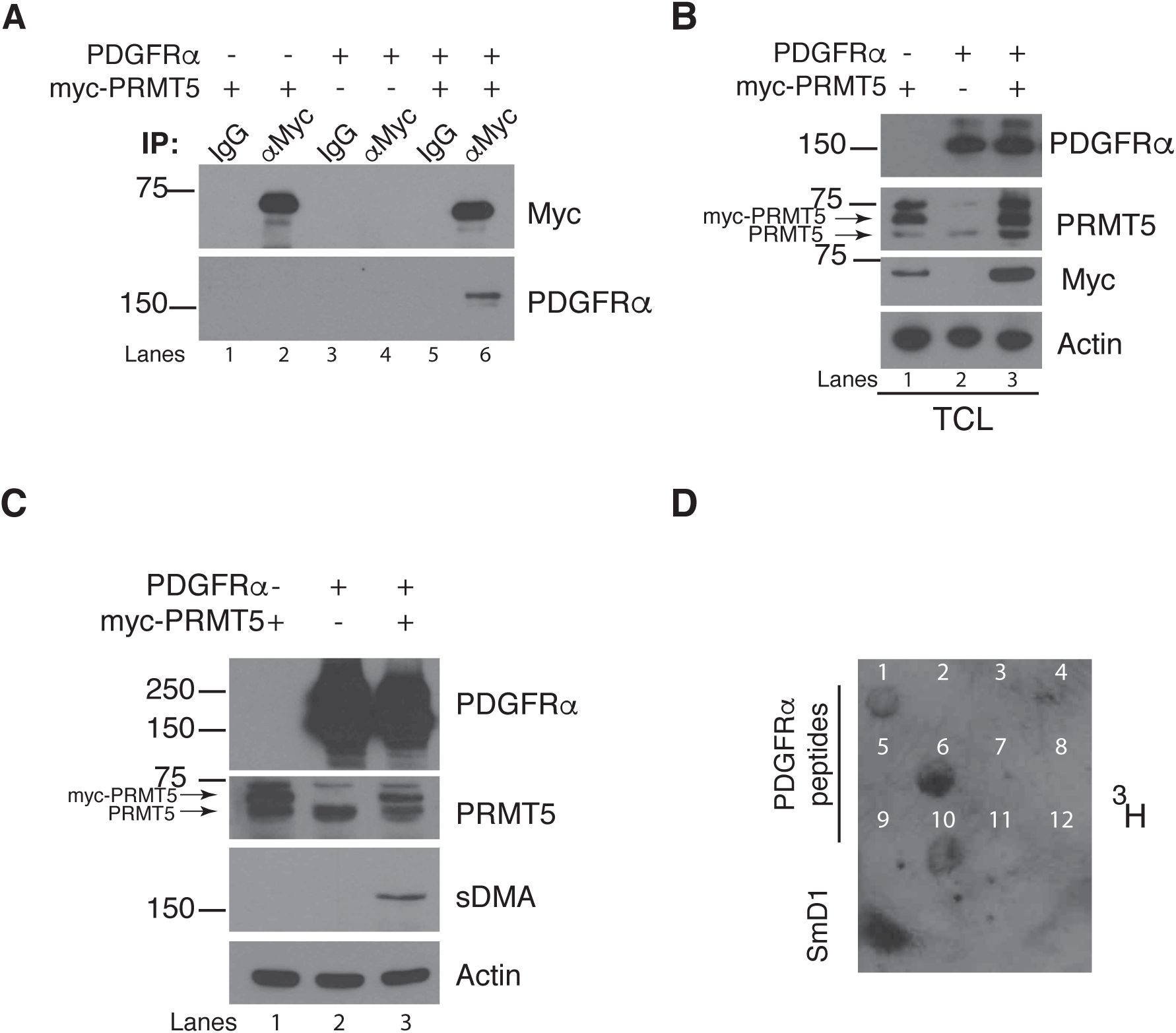
PDGFRα associates and is methylated by PRMT5. **A.** HEK293T cells transfected with myc-PRMT5 alone (lanes 1, 2), PDGFRα alone (lanes 3, 4) or myc-PRMT5 and PDGFRα (lanes 5, 6) were lysed an immunoprecipitated with either control IgG or anti-Myc antibody. The bound proteins were separated by SDS-PAGE and immunoblotted with anti-PDGFRα antibodies. **B.** Shown are total cell lysates from the expression of myc-PRMT5 alone, PDGFRα alone, or myc-PRMT5 and PDGFRα of panel A demonstrating equivalent expression. β-actin was used as an immunoblotting loading control. The molecular mass markers are shown on the left in kDa. **C.** HEK293T cells transfected with myc-PRMT5 alone (lane 1) PDGFRα alone (lane 2) or myc-PRMT5 and PDGFRα (lane 3) were lysed and immunoblotted with anti-sDMA antibodies. The molecular mass markers are shown on the left in kDa. **D.** *In vitro* methylation assay with the indicated PDGFRα peptides, showing positivity for peptides 1, 6 and 10. Peptide with the sequence from SmD1 was used as a positive control (n=4).

To identify PRMT5 methylation sites, we designed 12 peptides spanning the PDGFRα intracellular domain that represent potential methylated arginines. We next performed an *in vitro* methylation assay incubating these peptides with recombinant PRMT5-MEP50. We observed the methylation of peptides 1, 6 and 10 spanning amino acids 553-568, 907-921 and 812-826, respectively (Figure 6D). These regions correspond to the juxtamembrane domain and two sites in the second region of the tyrosine kinase domain, respectively. These findings define the PDGFRα as a PRMT5-MEP50 substrate.

### PDGFRα R554 is required for PRMT5 regulation of receptor stability

We generated arginine to lysine substitutions (R554K, R558/560K, R914K, R817/22K) of the identified PRMT5 arginines (Figure 6D) and examined their ability to regulate PDGFRα half-life. These PDGFRα:R-K proteins were expressed in HEK293T cells and their stability assessed by CHX treatment. Interestingly, PDGFRα:R554K had a decreased half-life in HEK293T cells without the need for PRMT5 depletion or inhibition, indicating that the loss of methylation at R554 is sufficient to decrease PDGFRα half-life (Figure 7A). Other R-K substitutions (PDGFRα:R558/60K, R914K, R817/22K) did not influence receptor half-life (Figure 7A).

**Figure 7.**
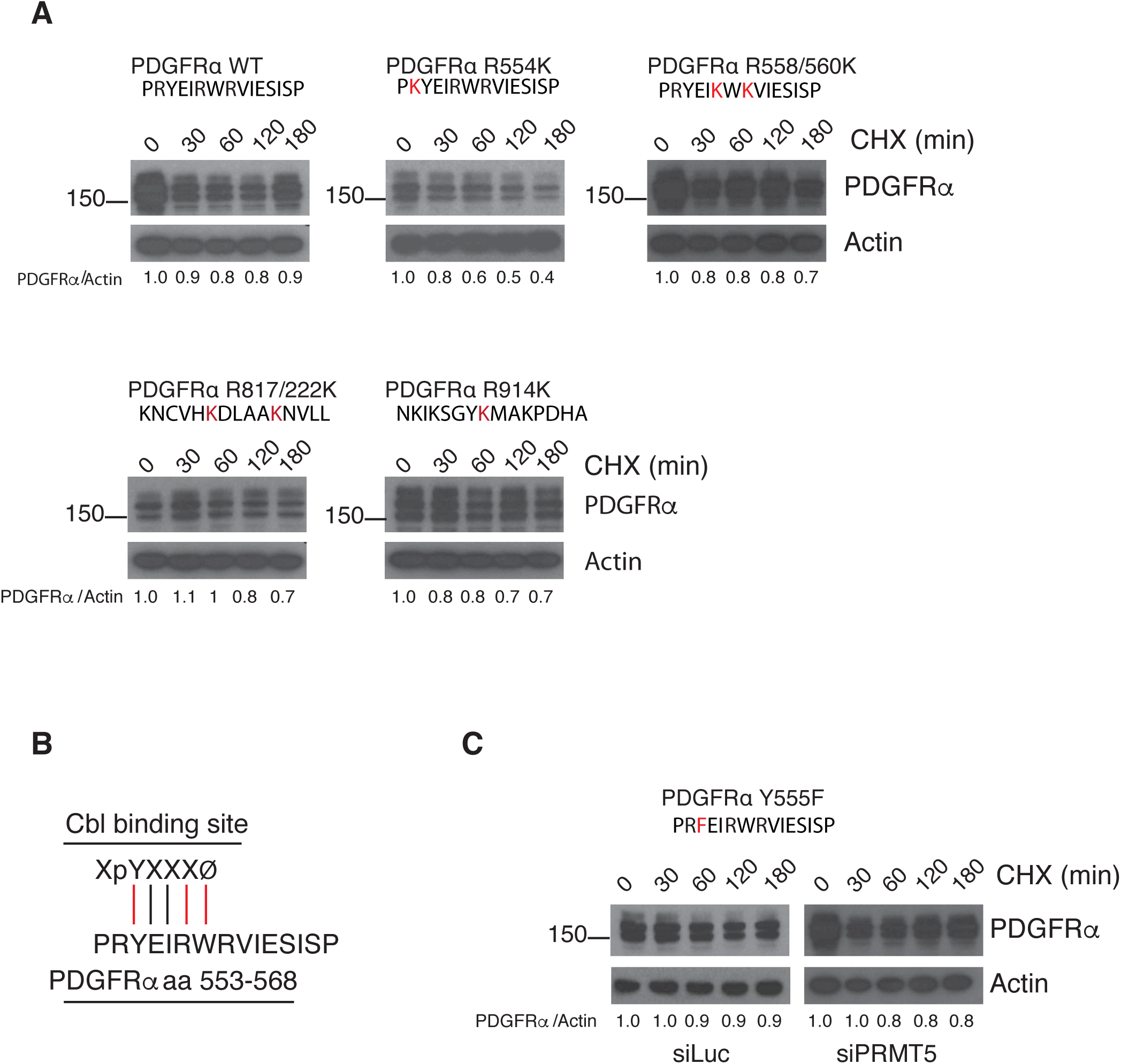
PRMT5 regulation of PDGFRα relies on R554. **A.** Western blot analysis of PDGFRα WT, R554K, R558/660K, R817/822K, and R914K mutants upon CHX treatment in 293T cells. CHX treatment was stopped at the time point indicated and PDGFRα protein levels were analysed by immunoblotting. β-actin was used as loading control. Relative PDGFRα/Actin ratio is shown and T0 was set as 1.0 (n=2 for R817/222K mutant, n=3 for others). **B.** Sequence of the human PDGFRα 553-568 peptide and the Cbl binding site. **C.** Western blot analysis of PDGFRα Y555F mutant upon CHX treatment in HEK293T cells transfected with siLuc or siPRMT5. CHX treatment was stopped at the time point indicated and PDGFRα protein levels were analysed by immunoblotting. β-actin was used as loading control. Relative PDGFRα/Actin ratio is shown, T0 was set as 1.0 (n=3).

We observed that the PDGFRα amino acids 553 to 558 harbours a potential Cbl binding site. Y555 is followed by a hydrophobic residue at pY+4 (W559, Figure 7B), in accord with the previous observation that determined XpYXXXØ as the Cbl consensus sequence (Lupher et al., 1997; Meng et al., 1999). Y555 is preceded by R554 (Figure 7B), therefore, we hypothesized that amino acids 553 to 558 of PDGFRα represents a Cbl binding site that is masked by arginine methylation. Thus, we substituted Y555F and assessed protein stability upon CHX treatment in HEK293T cells depleted of PRMT5. PDGFRα:Y555F was resistant to PDGFRα instability caused by PRMT5 deficiency (Figure 7C). Taken together, these data suggest that PRMT5 inhibition exposes a Cbl binding site (Y555) through the lack of R554 methylation.

## Discussion

In the present manuscript, we report that mice conditionally deleted for PRMT5 in OLs using Olig2-Cre have a severe hypomyelination phenotype that leads to death by the third post-natal week. OL precursor cells (PDGFRα+) were prevalent in brains of *PRMT5^FL/FL;Olig2Cre^* mice, eventhough there was a significant reduction in total number of OLs (Olig2+) including mature (CC1+) OLs. It is well-established that OPCs require signals from growth factors for self-renewal and proliferation. It is the time-controlled loss of these signals, in part, that initiates differentiation. We show that PRMT5 deficient OPCs display intracellular mislocalization of PDGFRα accompanied by decreased protein levels, thus impairing PDGFRα signaling capabilities to the downstream signaling pathways. Similar findings were observed using a specific PRMT5 inhibitor confirming the need for its methyltransferase activity. Mechanistically, PRMT5 associates with PDGFRα and methylates the receptor within a Cbl binding site (PDGFRα aa 553-568). Reduced PRMT5 activity leads to PDGFRα protein degradation in a Cbl-dependent manner. Our findings define a new role for PRMT5 as a regulator of receptor availability, representing an early signal to initiate OL differentiation. Accordingly, PRMT5 localization at the cell membrane has already been observed (Hsu et al., 2011). It has been shown that OPCs become unresponsive to the mitogenic stimulus of PDGF, by uncoupling the downstream signaling from the receptor (Hart et al., 1989), however, the mechanism has remained unknown. Our findings show that decreased PRMT5 expression during OL differentiation leads to trafficking defects promoting the ‘uncoupling’ of PDGF/PDGFRα with its signaling machinery. Indeed, in OPCs in which PRMT5 was depleted or inhibited by treatment with EPZ015666, we observed less PDGFRα OPC cell surface abundance and we observed alteration in PDGFRα distribution *in vivo* in the OPCs within the corpus callosum of *PRMT5^FL/FL;Olig2Cre^* mice. PDGFRα mislocalization correlates with decreased protein expression, thereby PRMT5 is involved in the regulation of the balance between recycling and degradation of the receptor *in vivo*. PRMT5 regulation of PDGFRα cellular homeostasis is required to ensure PDGFRα functionality and signaling, a balance that if altered, leads to defects in OPC homeostasis, differentiation and myelination. In OPCs, PRMT5 methylates PDGFRα maintaining its expression ensuring self-renewal and proliferation. During OL differentiation, PRMT5 expression decreases triggering a cascade of events required for proper differentiation, including PDGFRα downregulation.

PDGFRα trafficking and recycling is regulated by ubiquitination mediated by the Cbl E3 ubiquitin-protein ligase. Cbl regulates the availability of PDGFRα, where it triggers its ubiquitination and degradation (Bonita et al., 1997; Miyake et al., 1998). We show that PRMT5 contributes to the regulation of this process by controlling the availability of a Cbl binding site (PDGFRα aa 553-568). PRMT5 masks the availability of this site through arginine methylation of R554 thereby promoting the decreased half-life of PDGFRα in PRMT5-depleted cells (Figure 7C). This data are consistent with the observed progressive decrease of PRMT5 expression during OL differentiation (Figure 1E). Although our data define a new role for PRMT5, this function is accompanied by other known functions of PRMT5 in gene expression and pre-mRNA splicing regulation, as observed by our RNAseq analysis (Figure S2; Table S1). Thus it is evident that PRMT5 also fulfils nuclear roles that go beyond PDGFRα protein stability. Interplay between arginine methylation and phosphorylation has been observed before. Arginine methylation by PRMT1 interferes AKT phosphorylation of FOXO1 (Yamagata et al., 2008). Moreover, a crosstalk between R1175 and Y1173 has been observed, in which PRMT5 mediated methylation promotes tyrosine phosphorylation and ERK activation (Hsu et al., 2011).

Notably, a role for PRMT5 in the maintenance of pluripotency has already emerged. PRMT5 is required to promote pluripotency in the embryonic stem cells through epigenetic regulation (Tee et al., 2010). PRMT5 is also required for the homeostasis of precursor cells, including hematopoietic stem cells, neural stem/progenitor cells, skeletal muscle stem cells and chondrogenic progenitor cells ( Liu et al., 2011; Bezzi et al., 2013; Liu et al., 2015; Zhang et al., 2015; Norrie et al., 2016). PRMT5-mediated epigenetic regulation is also required to ensure adipogenic differentiation (LeBlanc et al., 2012).

In conclusion, our findings showing that PRMT5 inhibition causes a decrease in PDGFRα degradation in a Cbl-dependent manner may have therapeutic implications that go beyond OL differentiation. Inhibition of PRMT5 activity may regulate cell surface PDGFRα in certain cancers known to contain elevated PDGFRα expression such as glioblastomas (Ozawa et al., 2010; Puputti et al., 2006; Sun et al., 2014). Furthermore, as PRMT5 expression is low in mature OLs, the usage of PRMT5 inhibitors would likely not promote demyelination, although this remains to be shown.

## Author Contributions

S.C. and S.R. designed the experiments and wrote the paper. G.V. generated and maintained the mouse colony. S.C. and G.V. performed RNA extraction and qPCR, Western blot analyses. Z.Y. generated the PDGFRα mutants. L.D. obtained the brain slides; L.D. and S.C. performed the immunohistochemistry. K.C. and C.L.K. performed the RNA-seq analysis. T.B.N. shared reagents and unpublished data. S.C. conducted all the other experiments, analysed the results and performed statistical analyses. S.R. supervised the project.

## Acknowledgements

We thank Jasmin Coulombe-Huntington, Stéphane Laporte, Arthur Jacob, Daniel Garcia-Santos, Amel Hamdi and Prem Ponka for helpful discussions. We thank Christian Young for expert assistance with FACS analysis. This work was supported by grants from the Canadian Institute of Health Canada (MOP-93811 and PJT- 153203) to S. R., a salary award (C.L.K) and a doctoral fellowship (K.C.) from Fonds de recherche du Québec-Santé. Computational infrastructure was provided by Canada Foundation for Innovation (CFI) and by Compute Canada.

## STAR Methods

### Mouse strains and genotyping

All animal studies were approved by the McGill University Animal Care Committee. Embryonic stem cells engineered to harbour a conditional allele of PRMT5 with loxP and FRT sites flanking exon 7 of *PRMT5* were purchased from EUCOMM (Cat# : EPD0160_2). Briefly, two different clones were injected into C57BL/6 blastocysts to obtain chimeric animals. The chimeras were subsequently crossed with C57BL/6 mice and progeny assessed for germline transmission. To remove the neomycin resistance cassette, mice were then crossed with FLP recombinase expressing mice to promote recombination between FRT sites (Jax Strain#003946) resulting in a floxed allele (PRMT5*^FL^*). Genomic DNA was isolated from ear biopsies and a DNA fragment was amplified using the PRMT5 F1 and R1 primers. For the removal of *PRMT5* within the CNS, PRMT5^FL^ mice were crossed with Olig2-Cre transgenic mice. Verification of the PRMT5^-^ allele was assessed with the PRMT5 F1 and R3 primers.

### Immunohistochemistry

Brain immunohistochemistry was performed as previously described (Darbelli et al., 2016). Pictures were taken either using Zeiss LSM Pascal Confocal Imaging System or Confocal Wave FX SD. Antibodies used: Olig2 (Millipore), MBP (Sternberger Monoclonals), PDGFRα (Santa Cruz, Cell Signaling Technology), CC1 (Abcam, dilution 1:200), Alexa Fluor-488 or -546 IgG (Invitrogen, dilution of 1:200). For fluoromyelin, slides were incubated with fluoromyelin (Invitrogen) for 1 h at room temperature after blocking (dilution 1:300).

### Reverse transcription and quantitative PCR

Total RNA from tissues and cells were isolated in appropriate amounts of TRIzol (Invitrogen) according to manufacturer’s instruction. After digestion with DNase I (Promega), 4 μg of total RNA was retrotranscribed using M-MLV reverse transcriptase (Promega). Real-time PCRs were performed using PowerUp SYBR Mastermix (Life Technologies) on 7500 Fast Real-Time PCR System (Applied Biosystem). The efficiency of the primer used was tested according to the MIQE (Minimum Information for Publication of Quantitative Real-Time PCR Experiments) guidelines. Results were normalized as described in the figure legends using the ΔΔct method. All the primers used in this study are listed in Table S2.

### Plasmids

All the PDGFRα mutants were generated by 2-step PCR. HA-Cbl plasmid was from Moulay Alaoui-Jamali (McGill University).

### Protein extracts and immunoblot analysis

Brains were lysed in RIPA buffer (Sigma Aldrich) supplemented with protease inhibitors (Roche), incubated 30 minutes on ice then centrifuged at 12 000 rpm for 10 min at 4°C and supernatants were collected. Protein concentrations were determined using Bradford (Bio-Rad). Protein extracts were resolved by SDS-PAGE, transferred to a nitrocellulose membrane using an immunoblot TurboTransfer system (Bio-Rad), blocked for 1h at room temperature in TBS-T 5% milk and incubated with primary antibody, followed by incubation of secondary antibodies conjugated to horseradish peroxidase (Sigma Aldrich). Immunoblot signals were detected using chemiluminescence (Perkin Elmer). For detailed information about the primary antibodies used in this study see Supplemental Experimental Procedures.

### Cell culture, transfections and treatments

Primary rat and mouse OPC were isolated as previously described from newborn mice and Sprague Dawley rat brains (Almazan et al., 1993; Chen et al., 2007). For detailed information see Supplementary Experimental Procedures. For siRNA transfections, OPC were transfected with Lipofectamine RNAiMax (Invitrogen) according to manufacture’s instruction. siRNAs used are listed in Table S2. For PDGFRα activation analyses, OPC were starved in DMEM/F12 w/o PDGF-AA and bFGF for 2 h and then incubated with DMEM/F12 + PDGFAA for the time points indicated. HEK293T cells were grown in Dulbecco’s modified Eagle’s medium (Wisent), supplemented with 10% BCS, penicillin/streptomycin and sodium pyruvate and transfected with calcium phosphate with the indicated expression vectors. For PRMT5 inhibition, OPC were treated with EPZ015666 at 5μM concentration for 24h.

### Immunoprecipitation

For immunoprecipitations, 48 h after transfection cells were lysed in lysis buffer (1% Triton, 150 mM NaCl, 20 mM Tris-HCl pH 8.0, 100 mM sodium vanadate, 0.01% phenylmethanesulfonyl fluoride and protease inhibitors). Supernatants were collected and incubated with primary antibodies for 2 h on ice, then 40 μl of 50% Protein A-Sepharose slurry was added and incubated at 4°C for 1 h under constant rotation. The beads were then washed 4 times with lysis buffer, boiled with 40 μl of 2× Laemmli buffer and immunoblotted.

### *In vitro* methylation assay

PDGFRα peptides were synthesized by Top Peptide (Hillesborough, Tampa, FL). 10 μg of each peptide were spotted on a PVDF membrane, followed by incubation with 10 μl of [methyl-^3^H] S-adenosyl-L-methionine solution (15Ci/mmol stock solution, Perkin-Elmer) and 7 μg of PRMT5:MEP50 active complex (Sigma Aldrich) in methylation buffer (HEPES pH 7.4 500mM, NP40 0,01%, DTT 1 mM, PMSF 1 mM) for 3h at 37°C. PVDF membrane was then washed 3 times with TBS-T, treated with EN^3^HANCE (Perkin Elmer), according to manufacturer’s instructions and the reaction was visualized by fluorography.

### Flow cytometry analysis

For PDGFRα membrane localization, OPC were incubated with 1μg of PDGFRα (Cell Signaling Technology) antibody for 30 min at 4°C. Cells were then washed and incubated with Alexa-546 (Invitrogen) for 30 min at 4°C. After washing, cells were FACS sorted using a BD LSR Fortessa Analyser. Results were then analysed using FlowJo.

### RNA sequencing and analysis

RNA-sequencing (150bp paired-end) was performed using Illumina platform (NGS Core, Department of Epigenetics and Molecular Carcinogenesis, UT MD Anderson CCRD, Smithville, Tx). Sequencing runs were processed with Illumina CASAVA software. Trimmomatic v0.32 (Bolger et al., 2014) was used to trim reads, including removal of low-quality bases at the end of reads (phred33<30), clipping of the first three bases and clipping of Illumina adaptor sequences using the palindrome mode. We performed quality trimming with a sliding window, cutting once the average quality of a window of four bases fell below 30. We discarded reads shorter than 32 base pairs after trimming. Trimmed reads were aligned to the reference genome mm10 using STAR v2.3.0e (Dobin et al., 2013). Quality control was performed using metrics obtained with FASTQC v0.11.2, SAMtools (Li et al., 2009), BEDtools (Quinlan and Hall, 2010) and custom scripts. Bigwig tracks were produced with custom scripts, using BEDtools (Quinlan and Hall, 2010) and UCSC tools. Data were visualized using the Integrative Genomics Viewer (Thorvaldsdottir et al., 2013). For detailed information about the gene expression, alternative splicing and gene ontology analyses see Supplemental Experimental Procedures.

## REFERENCES

Almazan, G., Afar, D.E., and Bell, J.C. (1993). Phosphorylation and disruption of intermediate filament proteins in oligodendrocyte precursor cultures treated with calyculin A. J Neurosci Res. 36, 163–172.

Anders, S., Reyes, A., and Huber, W. Detecting differential usage of exons from RNA-seq data. Genome Res 22, 2008–2017 (2012).

Andreu-Perez, P., Esteve-Puig, R., de Torre-Minguela, C., Lopez-Fauqued, M., Bech-Serra, J.J., Tenbaum, S., Garcia-Trevijano, E.R., Canals, F., Merlino, G., Avila, M.A., and Recio, J.A. (2011). Protein arginine methyltransferase 5 regulates ERK1/2 signal transduction amplitude and cell fate through CRAF. Science Signaling 4, ra58.

Antonysamy, S., Bonday, Z., Campbell, R.M., Doyle, B., Druzina, Z., Gheyi, T., Han, B., Jungheim, L.N., Qian, Y., Rauch, C., et al. (2012). Crystal structure of the human PRMT5:MEP50 complex. Proc. Natl. Acad. Sci. USA 109, 17960–17965.

Armstrong, R.C., Harvath, L., and Dubois-Dalcq, M.E. (1990). Type 1 astrocytes and oligodendrocyte-type 2 astrocyte glial progenitors migrate toward distinct molecules. J. Neurosci Res. 27, 400–407.

Barres, B.A., Hart, I.K., Coles, H.S., Burne, J.F., Voyvodic, J.T., Richardson, W.D., and Raff, M.C. (1992). Cell death and control of cell survival in the oligodendrocyte lineage. Cell 70, 31–46.

Bedford, M.T., and Clarke, S.G. (2009). Protein arginine methylation in mammals: who, what, and why. Mol Cell 33, 1–13.

Bezzi, M., Teo, S.X., Muller, J., Mok, W.C., Sahu, S.K., Vardy, L.A., Bonday, Z.Q., and Guccione, E. (2013). Regulation of constitutive and alternative splicing by PRMT5 reveals a role for Mdm4 pre-mRNA in sensing defects in the spliceosomal machinery. Genes Dev. 27, 1903–1916.

Blanc, R.S., and Richard, S. (2017). Arginine Methylation: The Coming of Age. Mol Cell 65, 8–24.

Bolger, A.M., Lohse, M., and Usadel, B. (2014). Trimmomatic: a flexible trimmer for Illumina sequence data. Bioinformatics 30, 2114–2120.

Bonita, D.P., Miyake, S., Lupher, M.L., Jr., Langdon, W.Y., and Band, H. (1997). Phosphotyrosine binding domain-dependent upregulation of the platelet-derived growth factor receptor alpha signaling cascade by transforming mutants of Cbl: implications for Cbl’s function and oncogenicity. Mol. Cell. Biol. 17, 4597–4610.

Calver, A.R., Hall, A.C., Yu, W.P., Walsh, F.S., Heath, J.K., Betsholtz, C., and Richardson, W.D. (1998). Oligodendrocyte population dynamics and the role of PDGF in vivo. Neuron 20, 869–882.

Chen, Y., Balasubramaniyan, V., Peng, J., Hurlock, E.C., Tallquist, M., Li, J., and Lu, Q.R. (2007). Isolation and culture of rat and mouse oligodendrocyte precursor cells. Nature Protocols 2, 1044–1051.

Cho, E.C., Zheng, S., Munro, S., Liu, G., Carr, S.M., Moehlenbrink, J., Lu, Y.C., Stimson, L., Khan, O., Konietzny, R., et al. (2012). Arginine methylation controls growth regulation by E2F-1. EMBO J. 31, 1785–1797.

Darbelli, L., Vogel, G., Almazan, G., and Richard, S. (2016). Quaking regulates neurofascin 155 expression for myelin and axoglial junction maintenance. J. Neurosci. 36, 4106–4120.

Darbelli, L., Choquet, K. Richard, S. and Kleinman, C.L. (2017) Identification of alternative splicing events regulated by QKI in oligodendrocytes reveals self-splicing. Scientific Rep. (In press).

Dobin, A., Davis, C.A., Schlesinger, F., Drenkow, J., Zaleski, C., Jha, S., Batut, P., Chaisson, M., and Gingeras, T.R. (2013). STAR: ultrafast universal RNA-seq aligner. Bioinformatics 29, 15–21.

Eden, E., Navon, R., Steinfeld, I., Lipson, D., and Yakhini, Z. GOrilla: a tool for discovery and visualization of enriched GO terms in ranked gene lists. BMC Bioinformatics 10, 48 (2009).

Friesen, W.J., Paushkin, S., Wyce, A., Massenet, S., Pesiridis, G.S., Van Duyne, G., Rappsilber, J., Mann, M., and Dreyfuss, G.(2001). The methylosome, a 20S complex containing JBP1 and pICln, produces dimethylarginine-modified Sm proteins. Mol. Cell. Biol. 21, 8289–8300.

Fruttiger, M., Karlsson, L., Hall, A.C., Abramsson, A., Calver, A.R., Bostrom, H., Willetts, K., Bertold, C.H., Heath, J.K., Betsholtz, C., and Richardson, W.D. (1999). Defective oligodendrocyte development and severe hypomyelination in PDGF-A knockout mice. Development 126, 457–467.

Fyffe-Maricich, S.L., Karlo, J.C., Landreth, G.E., and Miller, R.H. (2011). The ERK2 mitogen-activated protein kinase regulates the timing of oligodendrocyte differentiation. J. Neurosci. 31, 843–850.

Hart, I.K., Richardson, W.D., Bolsover, S.R., and Raff, M.C. (1989). PDGF and intracellular signaling in the timing of oligodendrocyte differentiation. J. Cell Biol. 109, 3411–3417.

Hsu, J.M., Chen, C.T., Chou, C.K., Kuo, H.P., Li, L.Y., Lin, C.Y., Lee, H.J., Wang, Y.N., Liu, M., Liao, H.W., et al. (2011). Crosstalk between Arg 1175 methylation and Tyr 1173 phosphorylation negatively modulates EGFR-mediated ERK activation. Nat. Cell Biol. 13, 174–181.

Huang, J., Vogel, G., Yu, Z., Almazan, G., and Richard, S.(2011). Type II arginine methyltransferase PRMT5 regulates gene expression of inhibitors of differentiation/DNA binding Id2 and Id4 during glial cell differentiation. J. Biol. Chem. 286, 44424–44432.

Ishii, A., Fyffe-Maricich, S.L., Furusho, M., Miller, R.H., and Bansal, R.(2012). ERK1/ERK2 MAPK signaling is required to increase myelin thickness independent of oligodendrocyte differentiation and initiation of myelination. J. Neurosci. 32, 8855–8864.

Jansson, M., Durant, S.T., Cho, E.C., Sheahan, S., Edelmann, M., Kessler, B., and La Thangue, N.B. (2008). Arginine methylation regulates the p53 response. Nat. Cell Biol. 10, 1431–1439.

Kaushik, S., Liu, F., Veazey, K., Gao, G., Das, P., Neves, L., Li, K., Zhong, Y., Lu, Y., Giuliani, V., et al. (2017). Genetic deletion or small molecule inhibition of the arginine methyltransferase PRMT5 exhibit anti-tumoral activity in mouse models of MLL-rearranged AML. Leukemia Epub ahead of print, doi: 10.1038/leu.2017.206

Koh, C.M., Bezzi, M., Low, D.H., Ang, W.X., Teo, S.X., Gay, F.P., Al-Haddawi, M., Tan, S.Y., Osato, M., Sabo, A., et al. (2015). MYC regulates the core pre-mRNA splicing machinery as an essential step in lymphomagenesis. Nature 523, 96–100.

LeBlanc, S.E., Konda, S., Wu, Q., Hu, Y.J., Oslowski, C.M., Sif, S., and Imbalzano, A.N. (2012). Protein arginine methyltransferase 5 (Prmt5) promotes gene expression of peroxisome proliferator-activated receptor gamma2 (PPARgamma2) and its target genes during adipogenesis. Mol. Endo. 26, 583–597.

Li, H., Handsaker, B., Wysoker, A., Fennell, T., Ruan, J., Homer, N., Marth, G., Abecasis, G., Durbin, R., and Genome Project Data Processing, S. (2009). The Sequence Alignment/Map format and SAMtools. Bioinformatics 25, 2078–2079.

Li, Y., Chitnis, N., Nakagawa, H., Kita, Y., Natsugoe, S., Yang, Y., Li, Z., Wasik, M., Klein-Szanto, A.J., Rustgi, A.K., and Diehl, J.A. (2015). PRMT5 is required for lymphomagenesis triggered by multiple oncogenic drivers. Cancer Discov. 5, 288–303.

Liao, Y., Smyth, G. K., and Shi, W. featureCounts: an efficient general purpose program for assigning sequence reads to genomic features. Bioinformatics 30, 923–930 (2014).

Liu, F., Cheng, G., Hamard, P.J., Greenblatt, S., Wang, L., Man, N., Perna, F., Xu, H., Tadi, M., Luciani, L., and Nimer, S.D. (2015). Arginine methyltransferase PRMT5 is essential for sustaining normal adult hematopoiesis. J. Clin. Invest. 125, 3532–3544.

Liu, F., Zhao, X., Perna, F., Wang, L., Koppikar, P., Abdel-Wahab, O., Harr, M.W., Levine, R.L., Xu, H., Tefferi, A., et al. (2011). JAK2V617F-mediated phosphorylation of PRMT5 downregulates its methyltransferase activity and promotes myeloproliferation. Cancer Cell 19, 283–294.

Love, M. I., Huber, W., and Anders, S. Moderated estimation of fold change and dispersion for RNA-seq data with DESeq2. Genome Biol 15, 550 (2014).

Lupher, M.L., Jr., Songyang, Z., Shoelson, S.E., Cantley, L.C., and Band, H.(1997). The Cbl phosphotyrosine-binding domain selects a D(N/D)XpY motif and binds to the Tyr292 negative regulatory phosphorylation site of ZAP-70. J. Biol. Chem. 272, 33140–33144.

McKinnon, R.D., Waldron, S., and Kiel, M.E. (2005). PDGF alpha-receptor signal strength controls an RTK rheostat that integrates phosphoinositol 3’-kinase and phospholipase Cgamma pathways during oligodendrocyte maturation. J. Neurosci. 25, 3499–3508.

Meng, W., Sawasdikosol, S., Burakoff, S.J., and Eck, M.J. (1999). Structure of the amino-terminal domain of Cbl complexed to its binding site on ZAP-70 kinase. Nature 398, 84–90.

Miyake, S., Lupher, M.L., Jr., Druker, B., and Band, H.(1998). The tyrosine kinase regulator Cbl enhances the ubiquitination and degradation of the platelet-derived growth factor receptor alpha. Proc. Natl. Acad. Sci. USA 95, 7927–7932.

Norrie, J.L., Li, Q., Co, S., Huang, B.L., Ding, D., Uy, J.C., Ji, Z., Mackem, S., Bedford, M.T., Galli, A., et al. (2016). PRMT5 is essential for the maintenance of chondrogenic progenitor cells in the limb bud. Development 143, 4608–4619.

Ozawa, T., Brennan, C.W., Wang, L., Squatrito, M., Sasayama, T., Nakada, M., Huse, J.T., Pedraza, A., Utsuki, S., Yasui, Y., et al. (2010). PDGFRA gene rearrangements are frequent genetic events in PDGFRA-amplified glioblastomas. Genes Dev. 24, 2205–2218.

Pal, S., Vishwanath, S.N., Erdjument-Bromage, H., Tempst, P., and Sif, S.(2004). Human SWI/SNF-associated PRMT5 methylates histone H3 arginine 8 and negatively regulates expression of ST7 and NM23 tumor suppressor genes. Mol. Cell. Biol. 24, 9630–9645.

Puputti, M., Tynninen, O., Sihto, H., Blom, T., Maenpaa, H., Isola, J., Paetau, A., Joensuu, H., and Nupponen, N.N. (2006). Amplification of KIT, PDGFRA, VEGFR2, and EGFR in gliomas. Molecular cancer research: MCR 4, 927–934.

Quinlan, A.R., and Hall, I.M. (2010). BEDTools: a flexible suite of utilities for comparing genomic features. Bioinformatics 26, 841–842.

Raff, M.C., Lillien, L.E., Richardson, W.D., Burne, J.F., and Noble, M.D. (1988). Platelet-derived growth factor from astrocytes drives the clock that times oligodendrocyte development in culture. Nature 333, 562–565.

Richardson, W.D., Pringle, N., Mosley, M.J., Westermark, B., and Dubois-Dalcq, M.(1988). A role for platelet-derived growth factor in normal gliogenesis in the central nervous system. Cell 53, 309–319.

Schuller, U., Heine, V.M., Mao, J., Kho, A.T., Dillon, A.K., Han, Y.G., Huillard, E., Sun, T., Ligon, A.H., Qian, Y., et al. (2008). Acquisition of granule neuron precursor identity is a critical determinant of progenitor cell competence to form Shh-induced medulloblastoma. Cancer Cell 14, 123–134.

Shen, S., Park, JW., Lu, ZX., Lin, L., Henry, MD., Wu, YN., Zhou, Q., and Xing, Y., rMATS: robust and flexible detection of differential alternative splicing from replicate RNA-Seq data. Proceedings of the National Academy of Sciences of the United States of America 111, E5593–5601 (2014).

Sun, Y., Zhang, W., Chen, D., Lv, Y., Zheng, J., Lilljebjorn, H., Ran, L., Bao, Z., Soneson, C., Sjogren, H.O., et al. (2014). A glioma classification scheme based on coexpression modules of EGFR and PDGFRA. Proc. Natl. Acad. Sci. USA 111, 3538–3543.

Suzuki, R., and Shimodaira, H.(2006). Pvclust: an R package for assessing the uncertainty in hierarchical clustering. Bioinformatics 22, 1540–1542.

Tee, W.W., Pardo, M., Theunissen, T.W., Yu, L., Choudhary, J.S., Hajkova, P., and Surani, M.A. (2010). Prmt5 is essential for early mouse development and acts in the cytoplasm to maintain ES cell pluripotency. Genes Dev. 24, 2772–2777.

Thorvaldsdottir, H., Robinson, J.T., and Mesirov, J.P. (2013). Integrative Genomics Viewer (IGV): high-performance genomics data visualization and exploration. Brief Bioinform. 14, 178–192.

Xiao, J., Ferner, A.H., Wong, A.W., Denham, M., Kilpatrick, T.J., and Murray, S.S. (2012). Extracellular signal-regulated kinase 1/2 signaling promotes oligodendrocyte myelination in vitro. J. Neurochem. 122, 1167–1180.

Yamagata, K., Daitoku, H., Takahashi, Y., Namiki, K., Hisatake, K., Kako, K., Mukai, H., Kasuya, Y., and Fukamizu, A.(2008). Arginine methylation of FOXO transcription factors inhibits their phosphorylation by Akt. Mol. Cell 32, 221–231.

Yeh, H.J., Ruit, K.G., Wang, Y.X., Parks, W.C., Snider, W.D., and Deuel, T.F. (1991). PDGF A-chain gene is expressed by mammalian neurons during development and in maturity. Cell 64, 209–216.

Zhang, T., Gunther, S., Looso, M., Kunne, C., Kruger, M., Kim, J., Zhou, Y., and Braun, T.(2015). Prmt5 is a regulator of muscle stem cell expansion in adult mice. Nat. Commun. 6, 7140.

Zhao, Q., Rank, G., Tan, Y.T., Li, H., Moritz, R.L., Simpson, R.J., Cerruti, L., Curtis, D.J., Patel, D.J., Allis, C.D., et al. (2009). PRMT5-mediated methylation of histone H4R3 recruits DNMT3A, coupling histone and DNA methylation in gene silencing. Nat. Struct. Mol. Biol. 16, 304–311.

